# Regulation of sexual differentiation is linked to invasion in malaria parasites

**DOI:** 10.1101/533877

**Authors:** Gabrielle A. Josling, Jarrett Venezia, Lindsey Orchard, Timothy J. Russell, Heather J. Painter, Manuel Llinás

## Abstract

In the malaria parasite *Plasmodium falciparum*, the switch from asexual multiplication to sexual differentiation into gametocytes is essential for transmission to mosquitos. One of the key determinants of sexual commitment is the transcription factor PfAP2-G, which has been proposed to orchestrate this crucial cell fate decision by driving expression of gametocyte genes. We show conclusively that PfAP2-G is a transcriptional activator of gametocyte genes and identify the earliest known markers expressed during commitment. Remarkably, we also find that in sexually committed cells, PfAP2-G is associated with the promoters of genes important for red blood cell invasion and activates them through its interactions with a second transcription factor. We thus demonstrate an intriguing transcriptional link between the apparently opposing processes of red blood cell invasion and gametocytogenesis that is coordinated by the master regulator PfAP2-G. This finding has important implications for the development of new anti-malarial drugs that block the invasion of red blood cells by sexually committed cells, thereby preventing parasite transmission.

## Introduction

Though progress is being made towards the eradication of malaria, the disease continues to be a major global public health burden. Nearly half the world’s population is at risk of malaria and the disease results in hundreds of millions of deaths each year^1^. The malaria parasite *Plasmodium falciparum* has a complex lifecycle that involves development in the mosquito as well as the liver and red blood cells of humans. The symptoms of the disease are associated only with the 48-hour intraerythrocytic cycle, during which the parasite undergoes repeated rounds of multiplication, egress, and re-invasion as it passes through the ring, trophozoite, and schizont stages. However, these asexual stages cannot be productively taken up by the mosquito and transmitted. Instead, the parasite must undergo sexual differentiation to form male and female gametocytes. Crucially, only a small proportion of cells in each asexual cycle will commit. Commitment to sexual development generally occurs prior to schizogony, such that every merozoite within a single schizont will either enter sexual development or continue asexual multiplication following egress and re-invasion – this is known as next cycle conversion and seems to be the canonical pathway^2^. However, it is also possible for commitment to occur in the very early ring stage resulting in parasites developing directly into gametocytes without first passing through schizogony (same cycle conversion)^3^. During the 10-12 day process of gametocytogenesis, the parasite undergoes dramatic morphological changes as it passes through five distinct stages (stages I-V)^4,5^. With the exception of stage V gametocytes, gametocytes are sequestered in host tissues (particularly the bone marrow^6,7^).

One critical regulator of sexual commitment is AP2-G, which is a member of the ApiAP2 family of DNA-binding proteins^8,9^. Forward and reverse genetics in *P. falciparum*, *P. berghei*, and *P. yoelii* has demonstrated that AP2-G is a master regulator of gametocytogenesis^8-11^. Cells that do not express *ap2-g* will continue asexual development, whereas those that do are able to commit and enter gametocytogenesis. In *P. falciparum*, *ap2-g* is not expressed in most cells because it is epigenetically silenced by heterochromatin protein 1 (HP1) and histone deacetylase 2 (Hda2)^12-14^. HP1-mediated silencing of *ap2-g* is a ubiquitous feature of its regulation as it has recently been demonstrated across a range of *Plasmodium* species^15^. In *P. falciparum*, the protein GDV1 is responsible for removing HP1 from the *ap2-g* locus and so is also an important positive regulator of commitment^16,17^. Although some aspects of the regulation of PfAP2-G itself have now been explored, relatively little is known about how PfAP2-G directs the sexual commitment transcriptional program and which genes are involved. Furthermore, although PfAP2-G evidently plays a key role in regulating the commitment phase, its role in later stages of gametocytogenesis is unknown.

PfAP2-G is able to bind its motif in the promoters of gametocyte genes *in vitro* and it is able to activate expression of reporter genes, suggesting that it acts as an activator of transcription^8^. A recent study using single cell RNA-seq (scRNA-seq) revealed that PfAP2-G+ schizonts are chiefly characterised by a small number of up-regulated transcripts and very few that are down-regulated relative to asexual schizonts, which is consistent with this model^18^. Although collectively these data suggest that PfAP2-G likely directs commitment by binding the promoters of gametocyte genes and activating transcription, studies to date do not distinguish between direct and indirect effects of PfAP2-G *in vivo*. In addition, recent work indicates PfAP2-G may play a role in directing transcription beyond commitment, as it is present in the nucleus until stage I of gametocyte development^3^. Identification of the direct targets of PfAP2-G is thus crucial to understanding how commitment is regulated and will provide insight into the earliest events occurring during gametocytogenesis and define markers of sexual commitment.

In this study, we use the complementary approaches of chromatin immunoprecipitation followed by next generation sequencing (ChIP-seq) and transcriptomic characterization of a conditional knockdown cell line to identify targets of the transcription factor PfAP2-G. We find that PfAP2-G is not only associated with many known and novel early gametocyte promoters in the committed schizont phase, but also in sexual rings and stage I gametocytes. In addition, we demonstrate that PfAP2-G binding causes an increase in transcript levels by using CRISPR/Cas9 to mutate putative binding sites upstream of *ap2-g* itself. Surprisingly, we find that PfAP2-G is also associated with the promoters of a number of genes that are primarily associated with asexual development. These genes encode proteins involved in invasion of red blood cells. Many of these genes are also targets of the ApiAP2 protein PfAP2-I, a known transcriptional activator^19^. The two transcription factors not only have many common targets, but they also directly interact. We propose a model in which PfAP2-G has two major roles: it directs transcription of gametocyte genes to drive gametocytogenesis, and also interacts with PfAP2-I to cause an increase in invasion transcript levels.

## Results

### PfAP2-G regulates the expression of many early gametocyte genes

To examine global transcriptional changes associated with PfAP2-G activation, we used an AP2-G-DD parasite line^8^ in which PfAP2-G is endogenously tagged with an FKBP destabilisation domain (ddFKBP) and so can be conditionally stabilised or degraded depending upon the presence or absence of the small molecule Shld1 (Fig. 1a). This line produces gametocytes only in the presence of Shld1, though as in wildtype parasites only a subpopulation of cells commit in each cycle^8^. For the mRNA expression timecourse, we added Shld1 to AP2-G-DD parasites at the early trophozoite stage (24 hpi) and then collected RNA at six-hourly intervals for 66 hours for genome-wide transcriptomic analysis. This time period covers the development of committed schizonts into sexual rings and up to stage I gametocytes. We identified 42 genes that were differentially expressed as compared to a parallel culture grown in the absence of Shld1 (FDR ≤ 0.1, fold change ≥ 1.5). Consistent with our hypothesis that PfAP2-G is a transcriptional activator and with previous data^18^, all differentially regulated genes were upregulated in the presence of PfAP2-G (Fig. 1b). Virtually all of these transcripts are known to be expressed in committed schizonts or early gametocytes^12,17,18,20^, including many well-established early gametocyte genes (such as *pfs16*, *pfg14.744*, and *pfg27*) as well as novel genes and others that have only recently been implicated as potential early gametocyte genes. Of note, only a small number of transcripts were highly induced in committed schizonts (including *ap2-g* itself and *msrp1*), whereas many more genes showed altered expression following re-invasion.

**Figure 1:**
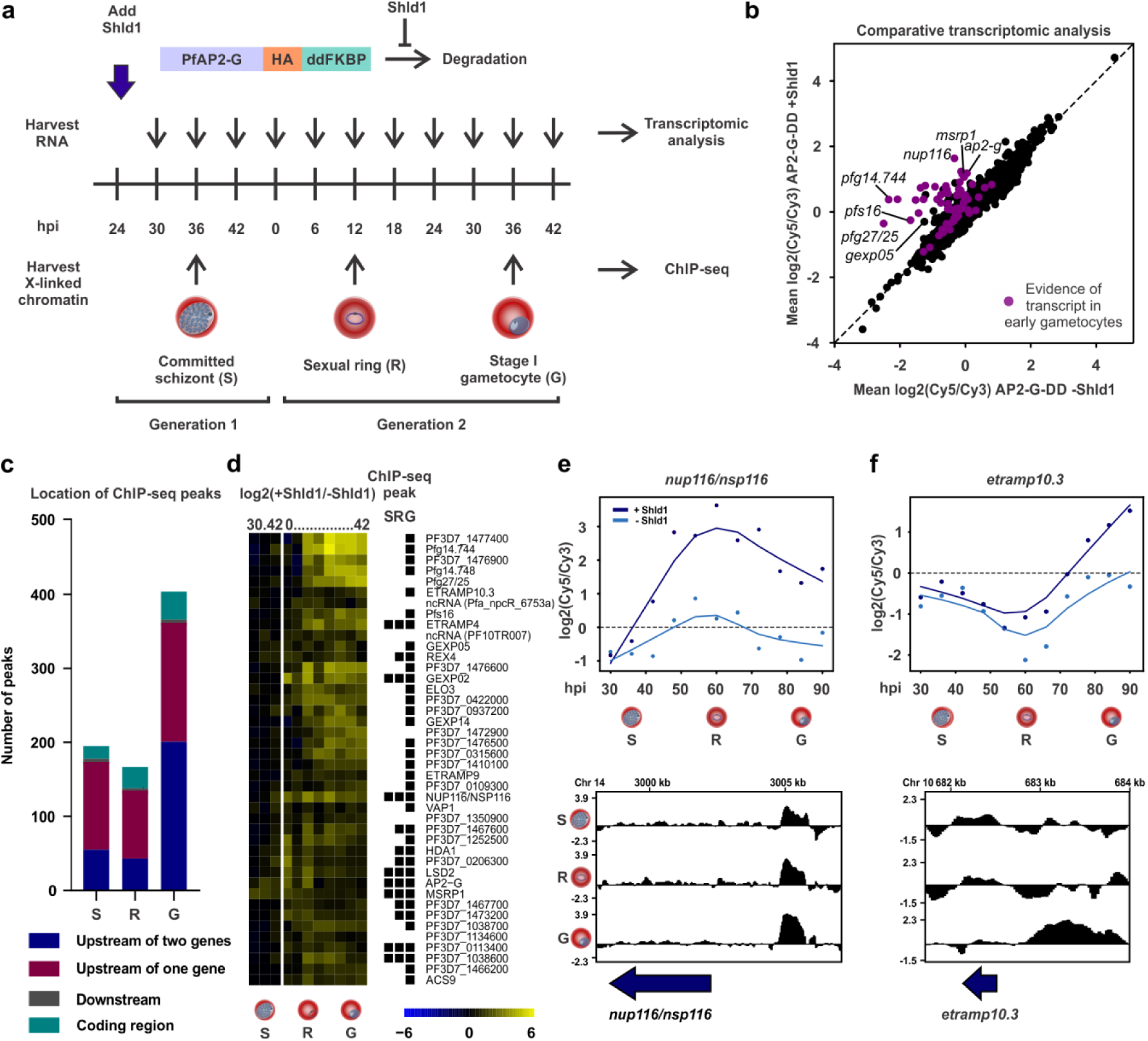
PfAP2-G binds the promoters of early gametocyte genes and its presence is associated with expression. **a**, Schematic showing the design of the transcriptomic and ChIP-seq experiments using AP2-G-DD. The AP2-G-DD line^8^, which allows for conditional stabilisation or degradation of PfAP2-G depending on the presence of Shld1, was used to identify PfAP2-G target genes. Transcript levels were compared in parallel cultures grown in the presence and absence of Shld1 and ChIP-seq was performed to identify PfAP2-G binding sites across the genome. Both experiments included committed schizonts (S), sexual rings (R), and stage I gametocytes (G). **b**, Scatterplot showing mean expression of all transcripts in AP2-G-DD +Shld1 versus AP2-G-DD -Shld1. Transcripts that have been identified in committed schizonts and early gametocytes^12,17,18,20^ are in purple. Several genes of particular interest are highlighted. **c**, Bar plot showing the number of PfAP2-G binding sites identified in each stage using ChIP-seq, with the location of each peak indicated by color. **d**, Heatmap showing expression of significantly differentially expressed transcripts (FDR ≤ 0.1 using Significance Analysis of Microarrays, fold change ≥ 1.5) over time in AP2-G-DD +Shld1 compared to AP2-G-DD -Shld1. The black squares to the right indicate whether or not the gene has a ChIP-seq peak upstream of it in each of the three stages tested. **e** and **f**, Expression of two PfAP2-G target genes (*nup116/nsp116* and *etramp10.3*) and PfAP2-G binding over time. The top plots show expression of the gene of interest in AP2-G-DD +Shld1 (dark blue) and AP2-G-DD -Shld1 (light blue). The bottom panels show log2-transformed PfAP2-G ChIP/input ratio tracks for committed schizonts, sexual rings, and stage I gametocytes at the locus of interest. The blue arrow shows the coding sequence.

We also performed chromatin immunoprecipitation followed by next generation sequencing in the same line to identify sites in the genome that are directly bound by PfAP2-G. ChIP-seq was performed at three stages that covered the same time period as the expression timecourse: committed schizonts, sexual rings, and stage I gametocytes. Because parasites were grown in the presence of Shld1 for the entire duration of the experiment, cells could potentially commit either at the end of cycle 1, the end of cycle 2, or the beginning of cycle 2 (if parasites undergo same-cycle conversion as recently described^3^). For simplicity we refer to parasites collected at the end of the second cycle as stage I gametocytes, though it is possible that this population consists of a combination of committed schizonts and stage I gametocytes produced via both same-cycle conversion and next-cycle conversion.

We identified 195 PfAP2-G binding sites in schizonts, 167 in rings, and 404 in stage I gametocytes (Fig. 1c). The vast majority of peaks for each stage were in intergenic regions, and most fell upstream of at least one gene. These binding sites were highly enriched in a DNA sequence motif very similar to that bound by PfAP2-G *in vitro*^21^ in all three stages, confirming the success of the ChIP (Supplementary Fig. 1). Importantly, almost all of the genes identified as possible PfAP2-G targets through the expression timecourse were also associated with PfAP2-G binding in at least one stage (Fig. 1d). The timing of PfAP2-G binding at the promoter of a gene generally corresponded well with when an increase in levels of the transcript under its control occurred (Fig. 1e, f). We also find that PfAP2-G clearly binds its own promoter from committed schizonts into stage I gametocytes and thus potentially auto-regulates its own expression. Overall, these data demonstrate that PfAP2-G is a direct activator of early gametocyte genes and thereby acts as the key driver of gametocytogenesis. PfAP2-G not only activates genes during the commitment phase but also up until at least stage I of gametocytogenesis. While this finding does not exclude a role for other transcriptional regulators during early gametocytogenesis, it does demonstrate that PfAP2-G continues to play a critical role in directing gametocytogenesis beyond commitment.

A number of potential downstream transcriptional regulators were identified in the transcriptomic analysis, such as the histone deacetylase Hda1 (PF3D7_1472200), the recently-identified putative histone demethylase LSD2 (PF3D7_0801900)^18^ and another ApiAP2 (PF3D7_1350900 or AP2-O4 in *P. berghei*)^22^. Several more genes encoding possible transcriptional regulators (including 12 ApiAP2 proteins) were identified by ChIP-seq but did not show significant changes in transcript levels in our transcriptomic analysis. This discrepancy may be because transcripts from the subpopulation of PfAP2-G+ cells were diluted by those from the PfAP2-G-cells also present in the population analysed, making it difficult to identify potentially subtle changes. We also identified many genes encoding exported proteins and several encoding proteins involved in fatty acid and lipid metabolism using both approaches (Fig. 1d). These findings fit with the known importance of host cell remodelling and lipids in gametocyte development^23,24^. Surprisingly, PfAP2-G was also associated with the promoters of several upsB *var* genes that encode PfEMP1 proteins that are important in immune evasion and cytoadhesion, though no significant changes in expression of any individual *var* gene were detected.

Notably, *gexp05* (PF3D7_0936600) was identified as a target of PfAP2-G in both the ChIP-seq and transcriptomic analyses. GEXP05 is an early gametocyte marker and has previously been shown to be expressed even in a parasite line that does not express functional PfAP2-G and so is thought to be regulated independently of it^25^. Our data suggest that although PfAP2-G is not strictly required for expression of *gexp05*, its presence nonetheless leads to increased transcription of the gene. This indicates that some degree of transcriptional regulation of gametocyte genes may occur upstream of PfAP2-G, possibly in a pre-commitment step that is augmented once PfAP2-G is present. In line with this, a recent study showed that many gametocyte genes are transcribed even in the absence of functional PfAP2-G but their transcripts are not stabilised resulting in aborted gametocyte commitment^26^.

### PfAP2-G is a transcriptional activator

Given the potential auto-regulation of *pfap2-g* expression, we sought to demonstrate that binding of PfAP2-G to a promoter leads to an increase in transcript levels by using CRISPR/Cas9 to mutate several putative binding sites upstream of the *ap2-g* gene. The *ap2-g* promoter contains eight copies of the PfAP2-G motif spread over 1.3 kb (Fig. 2a). Although the resolution of ChIP-seq is not sufficient to determine which motifs PfAP2-G is binding, based on the position of the PfAP2-G peaks it is possible that all eight are bound. Work in both *P. berghei* and *P. falciparum* suggests that PfAP2-G is regulated in part through a positive feedback loop in which it is able to bind its own promoter and further increase the levels of *ap2-g* transcript^8,9,18^. Our ChIP-seq data demonstrate that PfAP2-G directly binds its own promoter, but the functional consequences of this binding are unclear. Therefore, we chose to mutate three motifs that are within 45 bp of each other (Fig. 2a, box) and are positioned in the middle of one of the ChIP-seq peaks. Following successful mutation of the motifs (Supplementary Fig. 2), we used ChIP-qPCR to confirm these mutations abolished binding of PfAP2-G to its promoter (Fig. 2b). We also observed reduced PfAP2-G binding to another region in the *ap2-g* promoter that was not mutated and to the promoter of a second PfAP2-G target gene (*nup116*/PF3D7_1473700). The reduced binding of PfAP2-G to these other sites is likely because PfAP2-G protein levels were reduced due to perturbation of the positive feedback loop (Supplementary Fig. 3).

**Figure 2:**
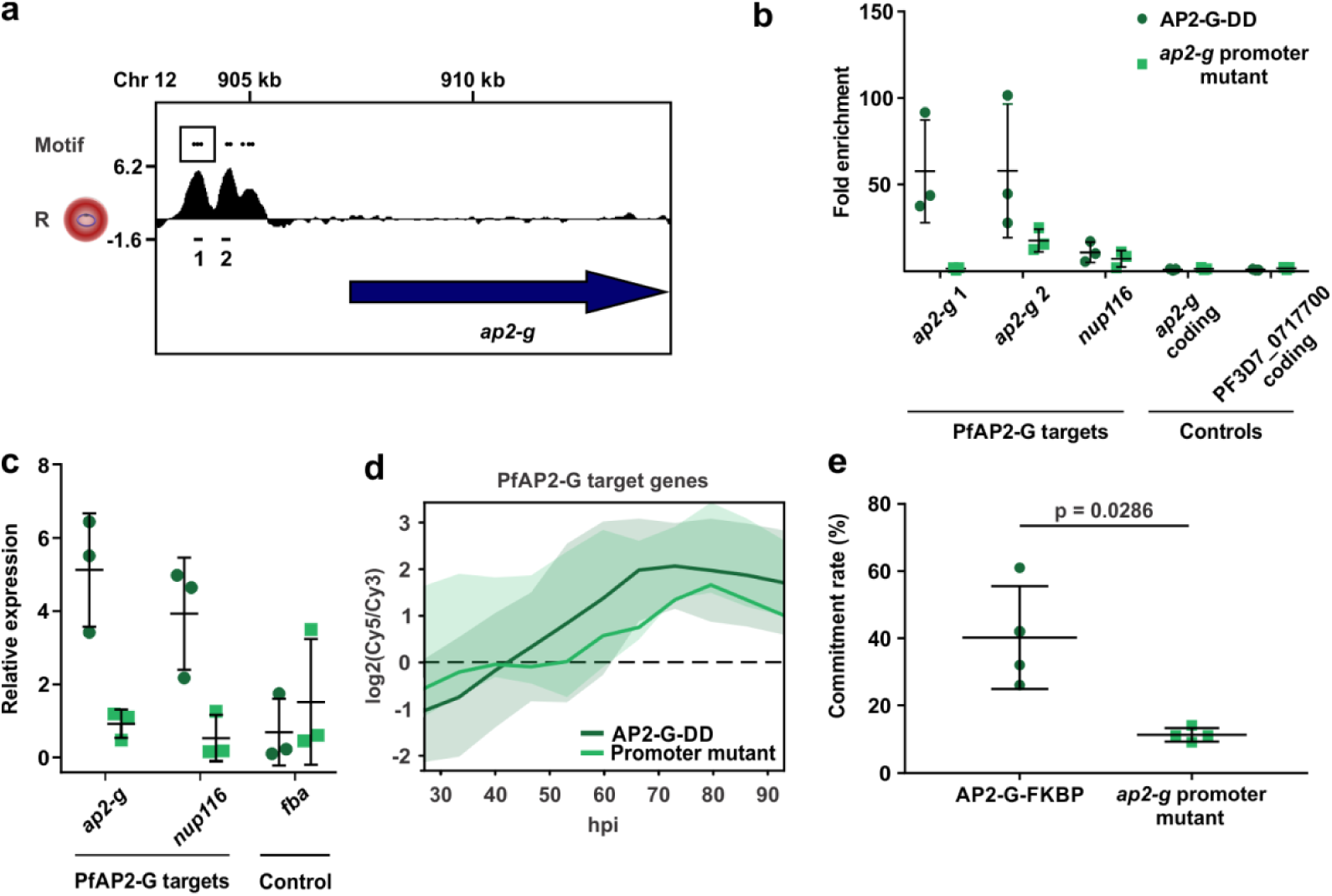
PfAP2-G regulates its own transcription. **a**, Log2-transformed PfAP2-G ChIP/input ratio tracks for the *ap2-g* locus in sexual rings. The blue arrow shows the coding sequence. The positions of the PfAP2-G motifs (GTACNC) in the promoter are shown above and the box highlights the three that were mutated. The black lines below indicate the regions of the promoter tested by qPCR in panel c (not to scale). **b**, ChIP-qPCR performed in ring-stage parasites shows that in the *ap2-g* promoter mutant, PfAP2-G is no longer able to bind the region of the promoter containing the mutations. It is still able to bind two other target sites, though at a reduced level compared to its parent. This is presumably due to lower levels of PfAP2-G protein in the *ap2-g* promoter mutant. The black bars indicate the mean and standard deviation. n = 3. **c**, qRT-PCR performed in ring-stage parasites shows that mutation of the *ap2-g* promoter leads to reduced levels of *ap2-g* transcript and that of a target gene. The black bars indicate the mean and standard deviation. n = 3. **d**, Plot showing mean temporal expression of the 42 known PfAP2-G targets (from Fig. 1d) in the *ap2-g* promoter mutant (light green) and the parental line (dark green). The shaded regions show the standard deviation. **e**, The *ap2-g* promoter mutant has a lower commitment rate than its parent. The horizontal bars indicate the mean and standard deviation. n = 4. The p-value was calculated using the two-tailed Mann-Whitney U test.

**Figure 3:**
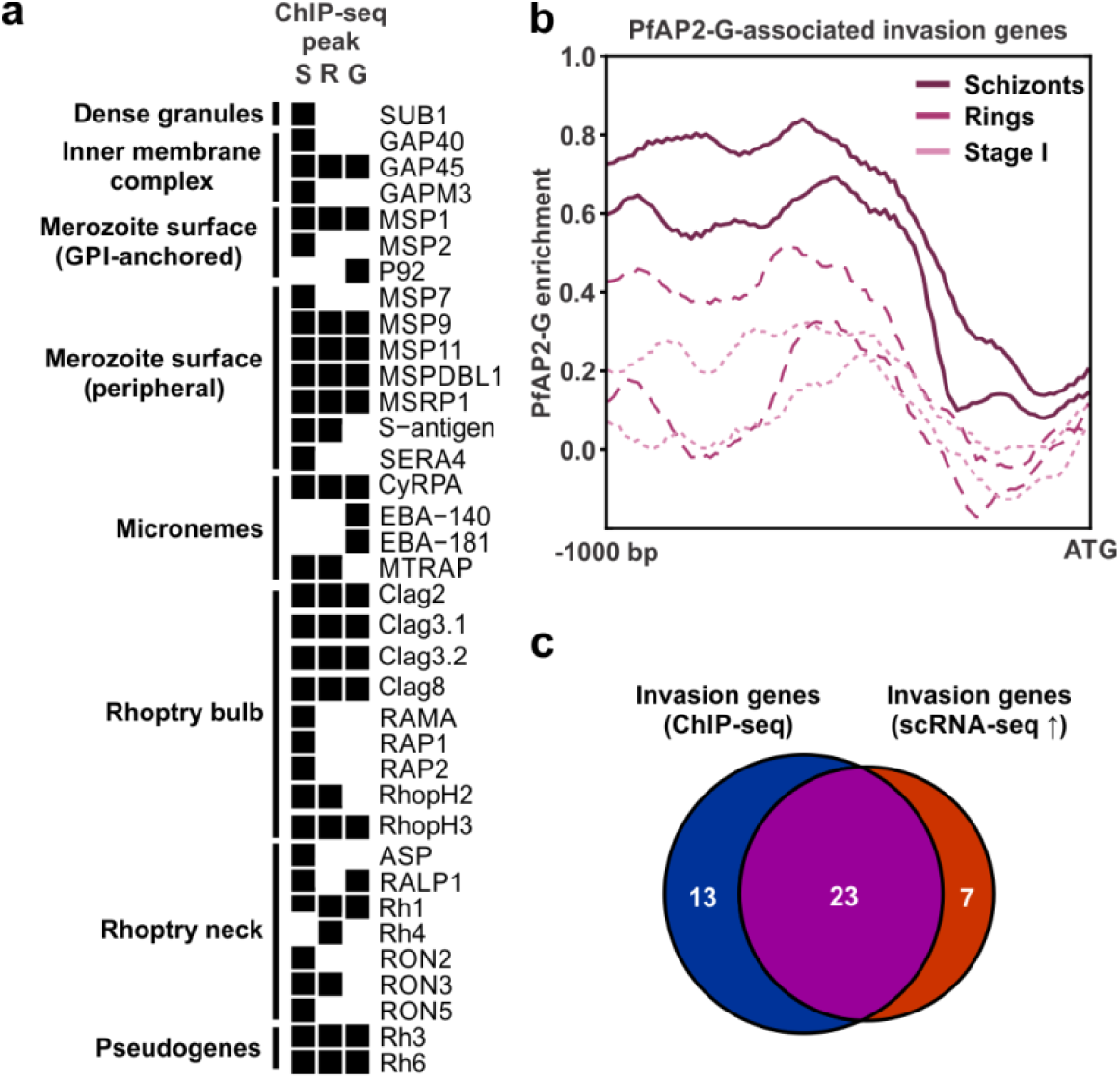
PfAP2-G binds the promoters of invasion genes. **a**, Invasion genes that are bound by PfAP2-G in one or more stage are listed on the left and are organized by compartment. The black squares to the right indicate whether or not the gene has a ChIP-seq peak upstream of it in each of the three stages tested. **b,** Plot showing the average enrichment of PfAP2-G across the 1000 bp upstream of the 36 invasion genes bound by PfAP2-G. Both biological replicates are shown for each stage, with enrichment in committed schizonts in dark purple, sexual rings in medium purple, and stage I gametocytes in pink. **c**, Venn diagram showing the overlap between invasion genes associated with PfAP2-G binding and those that were up-regulated in committed schizonts in either of two scRNA-seq studies^18,29^.

As expected, qRT-PCR showed that mutation of the *ap2-g* promoter led to reduced levels of *ap2-g* transcript and a target gene (*nup116*/PF3D7_1473700) in rings (Fig. 2c). To identify additional changes in transcript levels, genome-wide changes in mRNA abundance were measured. Overall, the *pfap2-g* promoter mutant did not display global changes in transcript levels or altered progression throughout the asexual cycle (Supplementary Fig. 4). Although *ap2-g* levels were not significantly reduced at all timepoints, they were down in the ring stage and overall the expression over time had a reduced range supporting a perturbation of *ap2-g* regulation (Supplementary Fig. 5). qRT-PCR performed at multiple stages confirmed that the down-regulation of *ap2-g* in the promoter mutant is restricted to early ring-stage parasites, highlighting this as the period when the auto-regulatory function of PfAP2-G occurs (Supplementary Fig. 5d). This reduction in *ap2-g* levels is evidently sufficient to interfere with PfAP2-G function, as PfAP2-G target genes identified in our initial transcriptional analysis (Fig. 1d) were down-regulated (on average by over 20%) in the *ap2-g* promoter mutant (Fig. 2d, Supplementary Fig. 5). The most significant changes occurred late in the timecourse when most PfAP2-G target genes are usually highly expressed, with 19 of the 42 genes significantly down-regulated in the final four timepoints (FDR ≤ 0.1, fold change ≤ 0.67). This down-regulation of PfAP2-G target genes results in a decrease in sexual commitment rate (Fig. 2e). Overall, these data establish that PfAP2-G indeed regulates itself through a positive feedback loop and confirm its functional role as a transcriptional activator of a number of early gametocyte genes.

### PfAP2-G regulates genes encoding proteins important for red blood cell invasion

Surprisingly, we also found that PfAP2-G is associated with the promoters of many genes encoding proteins with functions associated with red blood cell invasion (Fig. 3a). Invasion is an essential and extremely complex process involving many proteins that are located in multiple compartments of the cell (particularly the rhoptries, micronemes, and merozoite surface)^27^. PfAP2-G was enriched in the promoters of genes belonging to most of the major families of invasion genes: *msps*, *seras*, *rons*, *raps*, *ebas*, *rhs*, *gaps*, etc. This finding was unexpected, as invasion is a process that occurs specifically in the asexual stages and not in gametocytes. After a sexually-committed merozoite invades a red blood cell, the parasite undergoes gametocytogenesis within the same cell and thus does not need to re-invade. Although some invasion proteins also play a role in late gametocytes^28^, to our knowledge this is not true of most.

Intriguingly, although PfAP2-G is found in the promoters of invasion genes in all three stages, the majority of the binding and the highest degree of enrichment occurs in the schizont stage (Fig. 3b). On average these transcripts had slightly decreased abundance almost 48 hours after peak PfAP2-G binding (Supplementary Fig. 6), presumably reflecting the development of a subpopulation of cells (stage I gametocytes) that do not express invasion genes. The delay between binding of PfAP2-G and the down-regulation of invasion genes likely rules out a role for PfAP2-G as a direct repressor of invasion genes. In agreement with this, the majority of the invasion genes bound by PfAP2-G have recently been shown to be up-regulated in committed schizonts using scRNA-seq (Fig. 3c)^18,29^, suggesting that PfAP2-G likely enhances transcription of invasion genes.

Although PfAP2-G is associated with the promoters of many invasion genes, the only one that is strongly up-regulated in our transcriptomic analysis in the presence of PfAP2-G is *msrp1* (Fig. 1d, Supplementary Fig. 7a). MSRP1 has homology to MSP7, a known invasion protein, but little is known about it beyond its dispensability for asexual growth^30^. Multiple datasets show that *msrp1* is up-regulated in committed cells^16,18,29,31,32^. Crucially, *msrp1* is – like *ap2-g –* highly up-regulated in field isolates compared to laboratory strains, suggestive of a possible role in gametocytogenesis^33^. We tested a Δ*msrp1* line^30^ for its ability to form gametocytes to determine whether MSRP1 plays a role in gametocytogenesis or in invasion by committed merozoites. We found that this line produces gametocytes at a significantly higher rate than its parent (Supplementary Fig. 7b), suggesting that MSRP1 has a function in sexually-committed cells.

### PfAP2-G and PfAP2-I cooperate to regulate invasion genes

Many of the invasion genes bound by PfAP2-G are also targets of the transcription factor PfAP2-I^19^. PfAP2-I is another member of the ApiAP2 DNA-binding protein family that has recently been characterised as an activator of invasion genes^19^. Although ChIP-seq has been performed on PfAP2-I, comparing the PfAP2-G and PfAP2-I ChIP-seq datasets is difficult because these experiments were done in parasite lines with different genetic backgrounds (3D7 and Dd2, respectively). For this reason, we endogenously tagged PfAP2-I with GFP in the AP2-G-DD line and performed ChIP-seq on both proteins in parallel (Supplementary Fig. 8). We confirmed that the two proteins have many shared targets (Fig. 4a). Although many of the genes associated with binding of both proteins are invasion genes, there are also many that do not have roles in invasion (including three genes encoding ApiAP2 proteins) (Supplementary Fig. 9). The majority of these genes are down-regulated in response to stabilisation of PfAP2-G. Both proteins also have many unique targets; for example, the promoters of *sera6* and *sera7* are bound only by PfAP2-I (though PfAP2-G also binds *sera4*), whereas most early gametocyte genes (such as *pfg27, hda1*, and *nup116*) are only bound by PfAP2-G. PfAP2-I generally does not bind promoters of genes encoding micronemal or rhoptry neck proteins (as previously described^19^), but PfAP2-G does binds several of these (such as *eba175*, *ron2*, and *ron5*). Overall the summits of the ChIP-seq peaks for the two proteins are extremely close to each other, indicating that they are binding the same genomic locations (Fig. 4b). This suggests potential interplay between PfAP2-G and PfAP2-I.

**Figure 4:**
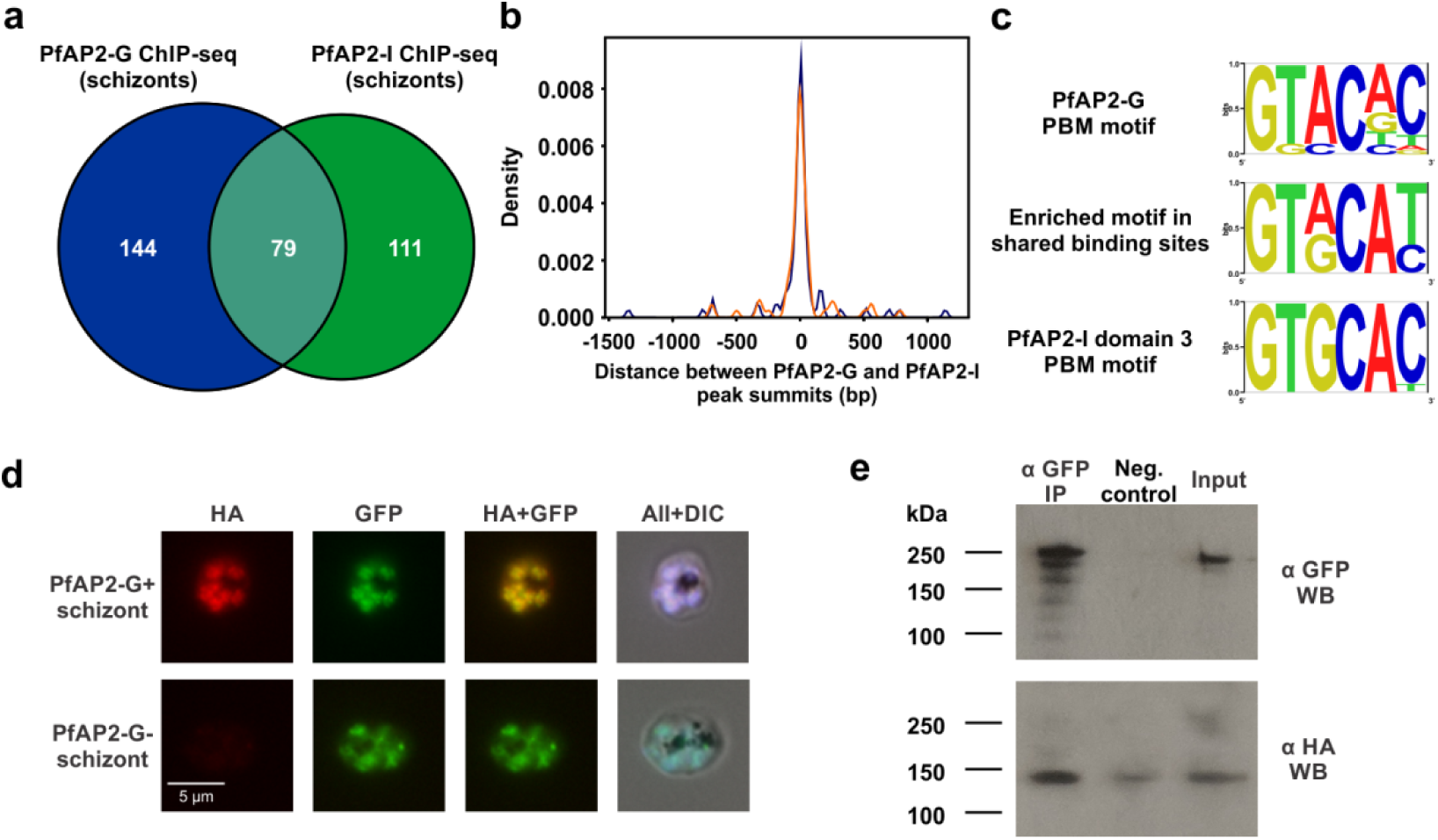
PfAP2-G interacts with the transcription factor PfAP2-I at invasion gene promoters. **a**, Venn diagram showing overlap between PfAP2-G and PfAP2-I binding sites in schizonts in ChIP-seq experiments performed using a parasite line expressing both AP2-G-HA-DD and AP2-I-GFP. **b**, Plot showing the distances between PfAP2-G and PfAP2-I peak summits within the 79 regions bound by both proteins. Replicate 1 is shown in blue and replicate 2 is shown in orange. **c**, The top enriched motif (determined by DREME) found in the regions bound by both PfAP2-G and PfAP2-I is a composite of the PfAP2-G and PfAP2-I protein-binding microarray DNA motifs. **d**, IFA images show that AP2-G+ cells also express PfAP2-I, though not all cells express PfAP2-G. **e**, Immunoprecipitation of AP2-I-GFP followed by Western blot with anti-HA and anti-GFP shows that PfAP2-I and PfAP2-G interact.

When we performed a motif analysis of regions of the genome bound by both PfAP2-G and PfAP2-I, the top result was a motif which is a composite of the PfAP2-G and PfAP2-I protein-binding microarray (PBM) motifs (Fig. 4c)^21^. The core PfAP2-G and PfAP2-I motifs are very similar: the PfAP2-G motif is GTACNC and the PfAP2-I motif is GTGCAC. The common motif had either a G or an A in the third position, giving a motif of GTRCAY. This indicates that PfAP2-G is not simply recruited by PfAP2-I and tethered to these regions indirectly, but is rather able to associate with at least some of these regions by directly binding DNA. Alternatively, interaction between the proteins may allow a minor change in DNA-binding preference of one of the two (or both).

To further investigate the potential interaction between PfAP2-G and PfAP2-I, we performed immunofluorescence assays. As previously demonstrated, PfAP2-G is not present in every cell^8^, but cells expressing PfAP2-G also express PfAP2-I (Fig. 4d). Immunoprecipitation experiments and ChIP-reChIP showed that PfAP2-G and PfAP2-I interact at invasion gene promoters (Fig. 4e, Supplementary Fig. 10). These findings indicate that PfAP2-G and PfAP2-I bind invasion promoters cooperatively to regulate transcription of invasion genes during commitment (Fig. 5), demonstrating that the regulation of gametocytogenesis and invasion are linked during malaria parasite development.

**Figure 5:**
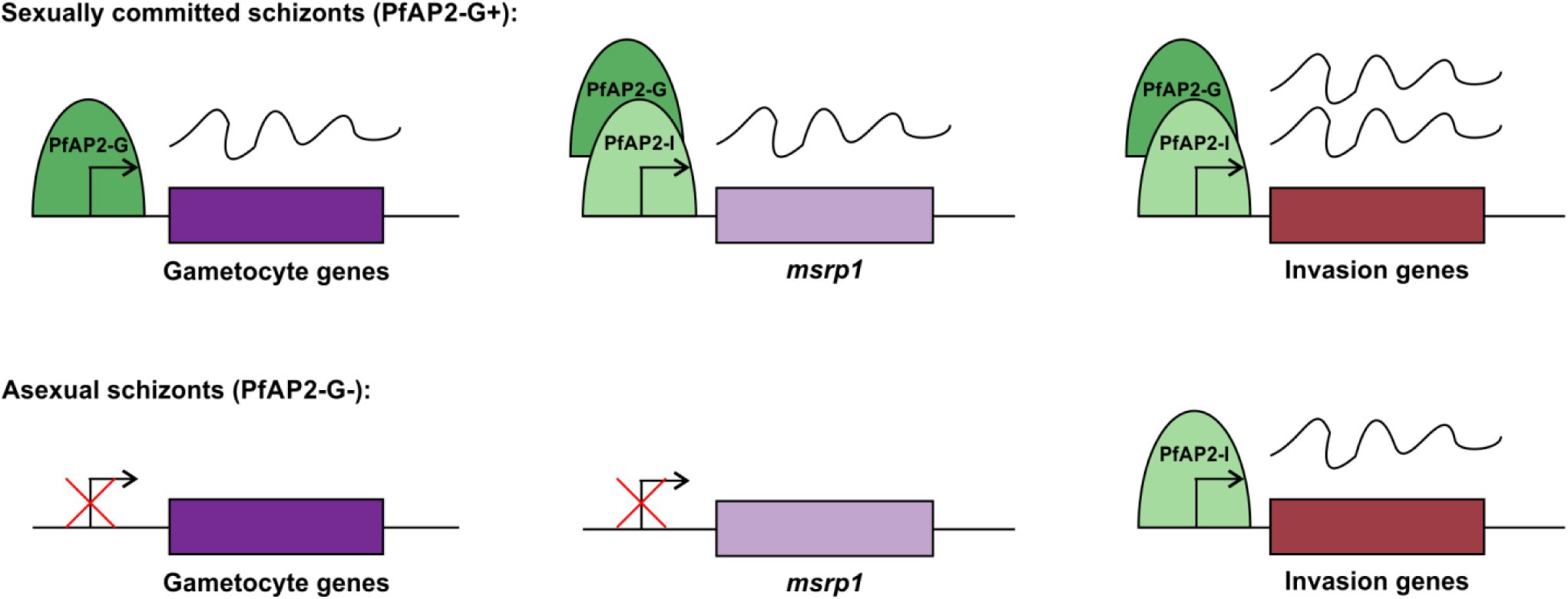
Model of regulation of gene expression by PfAP2-G during commitment. In sexually-committed schizonts, PfAP2-G is expressed and is able to bind the promoters of gametocyte genes and this leads to their expression. This binding thereby initiates the gametocyte transcriptional program. PfAP2-G also binds with PfAP2-I at invasion gene promoters (including *msrp1*) and this leads to increased levels of these transcripts. In contrast, in schizonts committed to continued asexual proliferation, PfAP2-G is not expressed and thus is unable to bind the promoters of early gametocyte genes and drive their expression. PfAP2-I is bound to the promoters of invasion genes where it activates transcription.

## Discussion

The master regulator AP2-G and the transcriptional program it directs are essential for sexual differentiation in the malaria parasite^8,9^. We have now identified targets of PfAP2-G and thus the earliest genes expressed during commitment, providing insight into the events occurring during early gametocyte development. Our results overall show that PfAP2-G positively regulates transcription of gametocyte genes, as has been previously hypothesized^8,9^. This is achieved in part by the regulation of PfAP2-G through a positive feedback loop in which it binds its own promoter to activate transcription, thereby generating high levels of PfAP2-G once the locus initially becomes active. Therefore, PfAP2-G establishes a transcriptional program that allows cells to irreversibly differentiate into the sexual stage of the parasite that is critical to mosquito transmission.^19,34^

Previous work has shown that PfAP2-G is present in the nucleus in committed schizonts^8^, so we expected to see PfAP2-G binding in this stage. However, it is noteworthy that PfAP2-G is also associated with the promoters of many genes in sexually committed rings and stage I gametocytes; in fact, PfAP2-G has many more binding sites in stage I gametocytes than in the other two stages. This indicates that PfAP2-G is not merely important during commitment, but also into the early stages of gametocyte development. The role of PfAP2-G in regulating gametocytogenesis thus extends beyond its essential role in directing the transcriptional switch from asexual multiplication to sexual differentiation. While this finding does not exclude a role for other transcriptional regulators during early gametocytogenesis, it does suggest that PfAP2-G likely continues to play a critical role in directing gametocytogenesis beyond commitment.

Surprisingly, we also found that PfAP2-G binds the promoters of invasion genes. We considered the possibility that PfAP2-G might be acting as a dual function transcription factor to repress invasion genes while activating gametocyte genes; however, this seems unlikely because although most invasion genes bound by PfAP2-G are subsequently down-regulated, this does not occur until 48 hours after peak PfAP2-G binding. As we present strong evidence that PfAP2-G activates expression of gametocyte genes, the most parsimonious explanation is that PfAP2-G also activates invasion genes in committed schizonts. In support of this hypothesis, two recent scRNA-seq studies have shown that many invasion genes are up-regulated in committed schizonts^18,29^. Another study that analysed transcript levels in fluorescently-sorted committed schizonts similarly showed that many invasion genes are slightly up-regulated when compared to asexual schizonts^32^. The fold changes observed in these studies were modest, so it is likely that slight increases in the levels of these transcripts were not detected by us due to the use of RNA from bulk cultures and the resulting dilution of signal from PfAP2-G-expressing cells.

Strikingly, many of the invasion genes regulated by PfAP2-G are also targets of the recently characterised regulator of invasion genes, PfAP2-I. Further, our data suggest that these factors interact in a way that is key to regulating invasion gene expression in the committed schizont. Combinatorial binding of transcription factors to the same promoters is an important mechanism of gene regulation that both increases specificity and allows finely-tuned regulation using a limited pool of transcription factors^35^. Many transcription factors are able to interact in such a way that increases their affinity for DNA, as in the case of enhanceosomes^36^. This may be the case for PfAP2-G and PfAP2-I, as AP2 domains have been shown to dimerize *in vitro*^37^ and in the related parasite *Toxoplasma gondii* cooperative binding of ApiAP2 proteins has been described^38^. An increase in DNA-binding affinity could explain why PfAP2-G binding leads to an increase in transcription of invasion genes. Apart from affecting affinity for DNA, the dimerization of transcription factors can lead to diverse outcomes, including a switch in function from repressing to activating (as in the case of Myc-Max and Mad-Max)^39^ and can even change the preferred DNA motif bound by a transcription factor in unpredictable ways^40^. This last possibility is particularly intriguing, as the PfAP2-G ChIP-seq motif only differs from its *in vitro* motif in the schizont stage, which is when PfAP2-I is also present (Supplementary Fig. 1). In future work, the identification of additional proteins that interact with PfAP2-G beyond PfAP2-I will be essential to clarifying how it regulates transcription. One possible interaction partner is PfBDP1, a bromodomain protein that is involved in activation of invasion genes and has been shown to interact with PfAP2-I^19,34^.

We propose a model in which PfAP2-G and PfAP2-I cooperate to direct a committed-schizont specific transcriptional program that includes up-regulation of many invasion genes (Fig. 5). In a committed schizont, many invasion transcripts are slightly up-regulated, with *msrp1* alone highly up-regulated during commitment across multiple datasets^12,16,18,29,31,32^. We demonstrate a role for MSRP1 in gametocytogenesis, as disruption of the gene leads to increased production of gametocytes. One explanation for this is that MSRP1 may be important for invasion of young red blood cells (reticulocytes) and hinder invasion of mature red blood cells. Committed cells grown only in the presence of mature red blood cells would thus be expected to be less likely to invade and subsequently develop into gametocytes, unless *msrp1* has been disrupted. In light of the significant up-regulation of *msrp1* that occurs in committed schizonts, the function of MSRP1 in committed cells requires further exploration.

One reason for this distinct invasion-related transcriptional program may be to allow committed merozoites to utilize a different invasion pathway than asexual merozoites. This could be analogous to the switch between sialic acid-dependent and -independent invasion pathways, which relies on altered expression of a very small number of genes^41^. A commitment-specific pathway may involve a switch in host cell tropism or reliance on different ligands to enter the red blood cell, or simply an improvement in invasion efficiency. In support of the first possibility, it has long been known that gametocytogenesis is increased when parasites are grown in the presence of reticulocytes^42,43^, and this is notable because of the presence of reticulocytes in the bone marrow where gametocytes accumulate^6,7^. In line with this, work in *P. berghei* has shown there is preferential formation of gametocytes in early reticulocytes within bone marrow^44^. The committed merozoite may have a preference for invasion of reticulocytes that is directed by PfAP2-G which could account for this propensity to develop in the bone marrow. This model is consistent with very recent data showing that a subset of *P. berghei* merozoites ‘home’ to the bone marrow where they develop within reticulocytes^45^. Importantly, this homing is receptor-mediated, indicating that there may be a transcriptional switch that allows this to occur. De Niz *et al*. similarly propose that committed merozoites may have a preference for invasion of cells in the bone marrow, and our work provides a possible transcriptional regulatory mechanism for this. If this model is correct, identifying the parasite ligands required for this preferential invasion will be an important next step.

In summary, our work shows definitively that the transcription factor PfAP2-G is a major cell fate determinant that triggers sexual commitment of the malaria parasite by activating genes required for early gametocytogenesis and that it continues to play a role beyond the commitment phase. Excitingly, we also show that PfAP2-G regulates invasion genes and thus connects the ostensibly disparate processes of gametocytogenesis and red blood cell invasion. Our observations support a model in which PfAP2-G directs a transcriptional program that allows committed merozoites to utilize a distinct invasion pathway. These insights significantly advance our knowledge of gametocyte biology, and may ultimately help in developing treatments that will prevent this crucial differentiation process and disrupt transmission of the malaria parasite.

## Methods

### Parasites and lines

Parasite cultures were maintained as previously described^19^. Synchronisation was performed using 5% sorbitol^46^. Parasites lines used were 3D7, AP2-G-DD^8^, AP2-G-DD^*ap2-g*^ ^*mut*^, AP2-G-DD::AP2-I-GFP, and 3D7Δ*msrp1*^30^. All AP2-G-DD-based lines were maintained in the presence of 2.5 nM WR99210 and routinely cultured in the absence of Shld1. 0.5 uM Shld1 was added at 24 hpi only when commitment to gametocytogenesis was required. The 3D7Δ*msrp1* line was maintained in the presence of 2.5 nM WR9910 except when commitment assays were performed. The AP2-G-DD^*ap2-g*^ ^*mut*^ line was created by transfecting uninfected red blood cells simultaneously with pUF-Cas9^47^ and pDC2-AP2-Gmut-BSD (described below) and adding AP2-G-DD trophozoites, as described^48^. The following day, cultures were fed using media containing 2.5 nM WR99210, 1 µg/mL blasticidin and 1.5 µM DSM1. To confirm successful mutation of the promoter, genomic DNA was purified using the Qiagen DNeasy Blood & Tissue kit and the region of interest was PCR amplified and Sanger sequenced. Parasites were then cultured only in the presence of WR99210 and cloned by limiting dilution. The AP2-G-DD::AP2-I-GFP line was made by transfecting the AP2-G-DD line with pSLI-AP2-I-GFP-BSD (described below) and selecting with 2.5 nM WR99210 and 1 µg/mL blasticidin. Once transfectants were obtained, they were selected for integration events using 400 µg/mL G418 as described^49^, genotyped by PCR, and then cloned by limiting dilution.

### Measurement of commitment rate

For AP2-G-DD-based lines, commitment was induced by addition of 0.5 uM Shld1 at the trophozoite stage (24 hpi). 3D7Δ*msrp1* and its parental line were grown for at least two weeks in the presence of 200 μM choline to suppress commitment^31^ and choline was removed only in trophozoite stage in the cycle the commitment assay began. For these experiments 3D7Δ*msrp1* was grown in the absence of WR99210. Cultures were treated with 50 mM N-acetyl glucosamine to kill asexual parasites for the four days following addition of Shld1 or the removal of choline, with daily media changes. The commitment rate was calculated as the gametocytemia 7 days after addition of Shld1 or removal of choline divided by the ring parasitemia the day following addition of Shld1. Four biological replicates were performed. Statistical comparisons were performed using the two-tailed Mann-Whitney U test.

### DNA constructs

To mutate the promoter of *ap2-g* using CRISPR/Cas9, complementary oligonucleotides encoding the guide RNA were inserted into pDC2-U6A-hDHFR (kind gift of Marcus Lee) using BbsI sites. A 633 bp homology region containing the three mutated PfAP2-G motifs (GTAC → GATC) was generated using overlap extension PCR, cloned into pGEM-T Easy (Promega), and then inserted using the NotI site. Finally, the *hDHFR* cassette was replaced with a *bsd* cassette obtained through PCR from pBcamHA using the NotI and SacI sites. The resulting plasmid was pDC2-AP2-Gmut-BSD.

The plasmid used to tag AP2-I with GFP in the AP2-G-DD line (pSLI-AP2-I-GFP-BSD) was modified from pSLI-TGD^49^. The *hDHFR* cassette was replaced with the *bsd* cassette from pBcamHA using the BamHI and HindIII sites and then a PCR product containing the 3’ end of *ap2-i* was inserted upstream of and in frame with *gfp* using the NotI and MluI sites.

### Chromatin immunoprecipitation

ChIP was performed as previously described^50^ with minor modifications. Shld1 was added to AP2-G-DD or AP2-G-DD::AP2-I-GFP cultures at 24 hpi to stabilize PfAP2-G and nuclei were harvested 12 (committed schizonts), 36 (sexual rings), or 60 (stage I gametocytes) hours later. Two biological replicates were used for ChIP-seq experiments and three for ChIP-qPCR. Crosslinked nuclei were then resuspended in Covaris shearing buffer (0.1% SDS, 10 mM Tris pH 8.1, 1 mM EDTA) and sonicated using an M220 ultrasonicator (Covaris) and the following conditions: 5% duty cycle, 75 W peak incident power, 200 cycles per burst, treatment time 300 seconds. The sonicated chromatin was diluted ten-fold with dilution buffer and precleared with Protein A/G magnetic beads (Pierce) for two hours. The pre-cleared material was then incubated with antibody and magnetic beads overnight, with a small volume of non-immunoprecipitated input material kept separately. For immunoprecipitations with AP2-G-DD, rat α HA (Roche 3F10) was used and for AP2-I-GFP rabbit α GFP (Abcam ab290) was used. For ChIP-qPCR, negative control IPs were also performed using either rat IgG (Abcam ab37361) or rabbit IgG (Abcam ab46540). The next day, beads were washed and the bound DNA eluted in elution buffer (1% SDS and 100 mM NaHCO_3_). Crosslinking was reversed by incubating overnight at 45°C and DNA was subsequently treated with RNase A at 37°C for 30 minutes and Proteinase K at 45°C for two hours. DNA was purified using the Qiagen MinElute kit and quantified using the Qubit HS DNA assay. ChIP DNA was either used to make libraries for ChIP-seq or analysed by qPCR as described below.

For ChIP-reChIP, the first ChIP was performed as described above up to the elution step. Beads were incubated with 10 mM DTT in TE for 30 minutes at 37°C and the recovered eluted DNA was then diluted 20-fold in dilution buffer. The diluted material was then subject to a second round of immunoprecipitation and eluted and processed as described above. 0.2 mg/mL HA peptide was included in the immunoprecipitation with α GFP to prevent re-precipitation with α HA still present.

### qPCR

All data were obtained using an Applied Biosystems 7300 Real-Time PCR machine and SDS v1.4 (Applied Biosystems). Data were analysed using the ΔΔCt method. All primer sets used for qPCR were first tested using serial dilutions of 3D7 genomic DNA to determine efficiency (primers were only used if >90%) and specificity (based on the presence of a single peak in the melting curve). For qRT-PCR, PF3D7_0717700 was used as a normalizing gene. For ChIP-qPCR, values are presented as fold enrichment in the immunoprecipitated sample versus the negative control (a mock IP performed with non-immune IgG). All qPCR experiments used two or three biological replicates.

### Library construction and analysis

Libraries for NGS sequencing were constructed as previously described^19^. Sequencing was performed using an Illumina Hiseq 2500 to generate 150 bp single-end reads.

### ChIP-seq data analysis

Following assessment of the quality of the reads using FastQC^51^, adapter sequences were trimmed using Trimmomatic^52^ and reads mapped to the *P. falciparum* 3D7 genome (Pf 3D7 v28, obtained from PlasmoDB) using BWA-MEM^53^. Reads were filtered using SAMtools^54^ to removes duplicates. For visualisation, bam files were converted to bigwig files showing log2(IP/input) using bamcompare from the deeptools suite^55^ and viewed using IGV^56^. MACS2^57^ was used to call peaks with a q value cutoff of 0.01. Called peaks were annotated using BEDtools^58^. Two biological replicates were used for each experiment and subsequent analysis focused on regions bound only in both samples (identified using the Multiple Intersect function of BEDtools). DREME^59^ was used to identify enriched DNA motifs in sequences associated with called peaks and TOMTOM^60^ was then used to compare any identified motifs with known ApiAP2 motifs^21^. Sequence logos were generated use enoLOGOS^61^. The plotProfile tool in the deeptools suite^55^ was used to plot the average enrichment of PfAP2-G in the 1000 bp upstream of the 36 invasion genes it is associated with.

For the generation of motif heatmaps, instances of the most enriched overall motifs for each dataset were identified within the overlapping peaks using FIMO^62^, with a significance threshold of 1e-3 against a second order Markov model. The top scoring motif was retained for each peak. A four color nucleotide plot was generated using the cegr-tools program FourColorPlot, available at https://github.com/seqcode/cegr-tools. Each nucleotide sequence was centered on the top scoring instance of the motif of interest, and a ten nucleotide extension was added in either direction of the conserved motif occurrence.

### RNA purification and cDNA synthesis

RNA was extracted from synchronised parasites by resuspending infected red blood cells in TRIzol (ThermoFisher Scientific) and following the manufacturer’s protocol. For qRT-PCR, cDNA was synthesized from 2 μg RNA using SuperScript II Reverse Transcriptase (ThermoFisher) with random nonamers and oligo(dT) primers. Negative controls without reverse transcriptase were also prepared to determine by qPCR whether contaminating gDNA was present.

### DNA microarrays

DNA microarrays were also performed as previously described^19,63^, except for the addition of cDNA from the samples to the normal asexual reference pool (to make up 50-63% of the new reference pool) to ensure that gametocyte-specific transcripts would be represented in the reference and thus could be quantified. Data were analysed using the Significance Analysis of Microarrays^64^ package in R. In scatter plots showing microarray data, the Lowess curve is shown in addition to the individual data points.

### Immunofluorescence assays

IFAs were performed as previously described^65^. PfAP2-G and PfAP2-I were detected with 1 μg/mL rat α HA (Roche 3F10) and 2 μg/mL rabbit α GFP (Abcam ab290), respectively. Secondary antibodies used were 2 μg/mL AlexaFluor 546 coupled goat α-rat (ThermoFisher Scientific) and 2 μg/mL AlexaFluor 488 coupled goat α-rabbit (ThermoFisher Scientific). Images were obtained using an Olympus BX61 system and SlideBook 5.0 (Intelligent Imaging Innovations).

### Immunoprecipitation

Immunoprecipitation experiments were performed using 36 hpi AP2-G-DD::AP2-I-GFP schizonts to which Shld1 had been added 12 hours earlier. Nuclear proteins were extracted using a conventional protocol^66^, and then diluted 1/4 in dilution buffer with 0.025% Tween-20 and added to washed GFP-Trap_MA beads (Chromotek). For the negative control, binding control magnetic agarose beads (Chromotek) were used. Following incubation at 4°C for one hour while rotating and five washes with PBS-T, bound proteins were eluted by boiling in loading buffer.

### Western blot

Proteins were run on a 4-12% Bis-Tris gel (ThermoFisher) in MES buffer and then transferred to a nitrocellulose membrane. After blocking with 5% milk in PBS-T for 30 minutes, membranes were incubated overnight with the primary antibody in 5% milk. Primary antibodies used were: 1 μg/mL rat α HA (Roche 3F10), 1/2000 rabbit α GFP (Abcam ab290), 1/1000 rabbit α aldolase HRP (Abcam ab38905), or 1/3000 mouse α H3 (Abcam ab10799). The next day, membranes were washed three times with PBS-T and incubated for two hours with the appropriate secondary antibody, then washed again. Secondary antibodies used were: 1/3000 goat anti-rat HRP conjugate (Millipore), 1/10 000 goat anti-rabbit HRP conjugate (Millipore), or 1/5000 goat anti-mouse HRP conjugate (Pierce). ECL reagent (Pierce) was used to detect bound antibody.

## Data availability

ChIP-seq data are deposited in the NCBI Sequence Read Archive (SRA) under the accession numbers GSE120448 (PfAP2-G enrichment in committed schizonts, sexual rings, and stage I gametocytes) and GSE120488 (PfAP2-G and PfAP2-I enrichment in schizonts). Microarray data are deposited in the NCBI Gene Expression Omnibus (GEO) under the accession numbers GSE120990 (AP2-G-DD + versus - Shld1) and GSE121312 (AP2-G-DD^*ap2-g mut*^ and AP2-G-DD).

## Acknowledgments

This work was funded through NIH grant R01 AI125565 and generous support from The Pennsylvania State University. G.A.J. is a recipient of the Sir Keith Murdoch Fellowship from the American Australian Association and a Postdoctoral Research Grant from the American Heart Association (16POST26420067).

## Author contributions

G.A.J. and M.L. designed the experiments. G.A.J., J.V., L.O., and H.P. performed the experiments. G.A.J. and T.J.R. analysed the data and generated figures. G.A.J. and M.L. wrote the manuscript.

## Additional information

Correspondence and requests for materials should be addressed to M.L.

