## Supplementary figures for "Regulation of sexual differentiation is linked to invasion in malaria parasites"

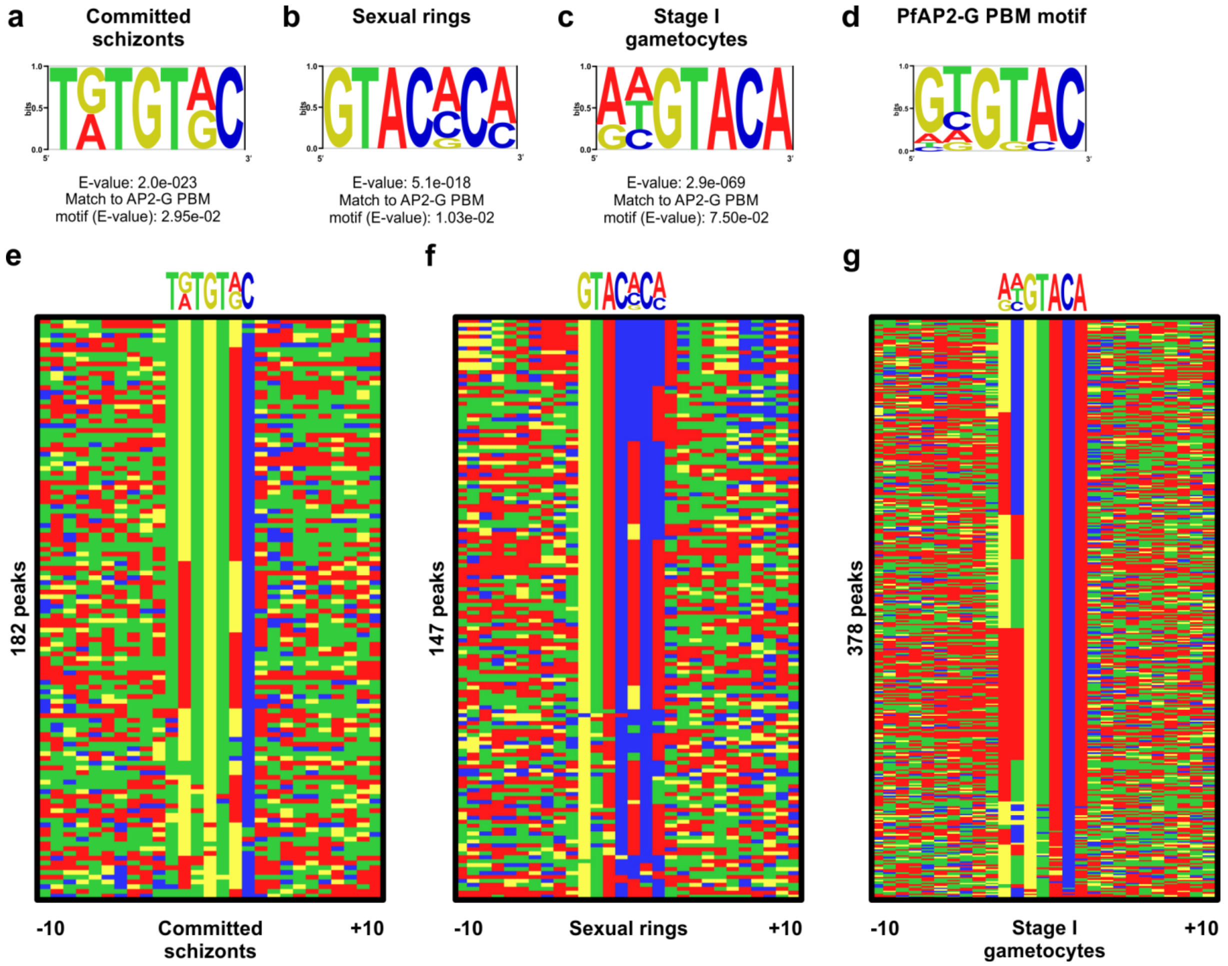


**Supplementary Figure 1:** DNA motif analysis of ChIP-seq peaks using DREME identified GTRC (**a**: committed schizonts) and GTAC (**b**: sexual rings and **c**: stage I gametocytes) as the top-scoring motifs. Comparison of each motif to known DNA motifs bound by recombinant AP2 domains using Tomtom showed that all three motifs match most closely to the DNA motif bound by the AP2 domain of AP2-G (**d**). DNA motif heatmaps of the ChIP-seq peaks and motifs from committed schizonts (**e**), sexual rings (**f**), and stage I gametocytes (**g**).

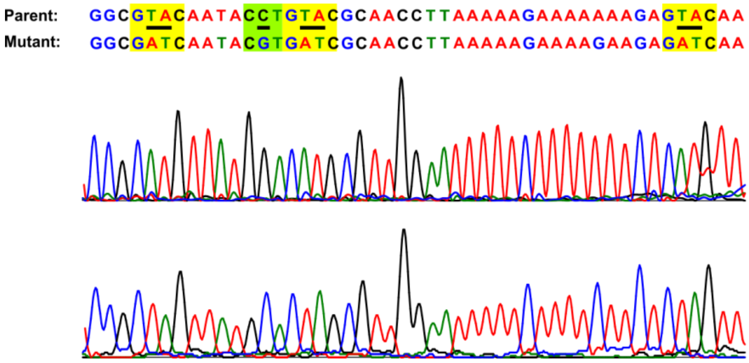


**Supplementary Figure 2:** DNA sequence chromatogram showing successful mutation of three of the eight PfAP2-G motifs upstream of *ap2-g*. The motifs are highlighted in yellow and the PAM is highlighted in green. Mutated nucleotides are underlined.

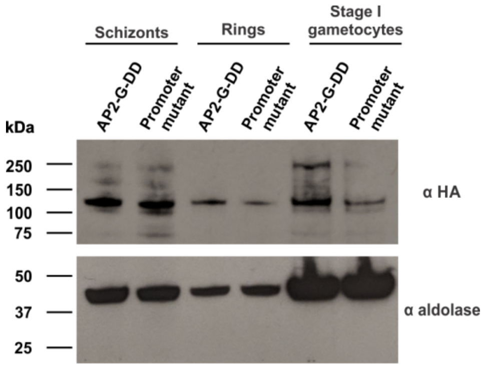


**Supplementary Figure 3:** Western blot showing PfAP2-G protein levels in the AP2-G-DD line and the *ap2-g* promoter mutant parasite line. Aldolase (bottom panel) was used as a loading control.

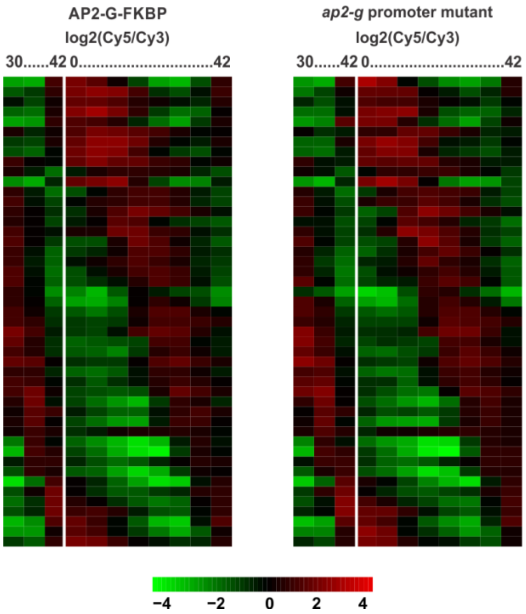


**Supplementary Figure 4:** Heatmaps showing expression of control genes[1] in AP2-G-FKBP and the *ap2-g* promoter mutant line. Both lines show a similar pattern of periodic gene expression, indicating that transcript levels are not globally affected and staging of the two lines is similar.

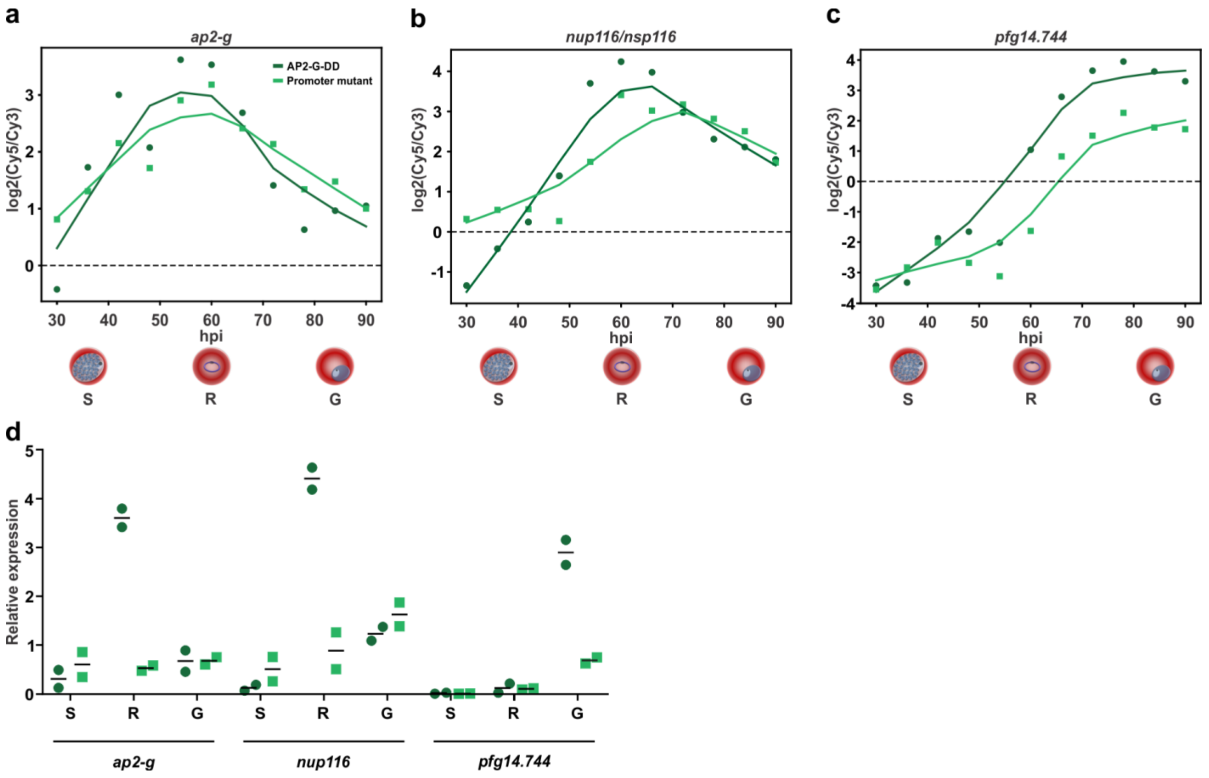


**Supplementary Figure 5:** Expression of **a**, *ap2-g*, **b**, *nup116/nsp116*, and **c**, *pfg14.744* in the *ap2-g* promoter mutant parasite line (light green) and its parent AP2-G-DD (dark green) in DNA microarrays. **d**, qRT-PCR analysis of RNA from three stages validates the DNA microarray data. The black bar indicates the mean. n = 2.

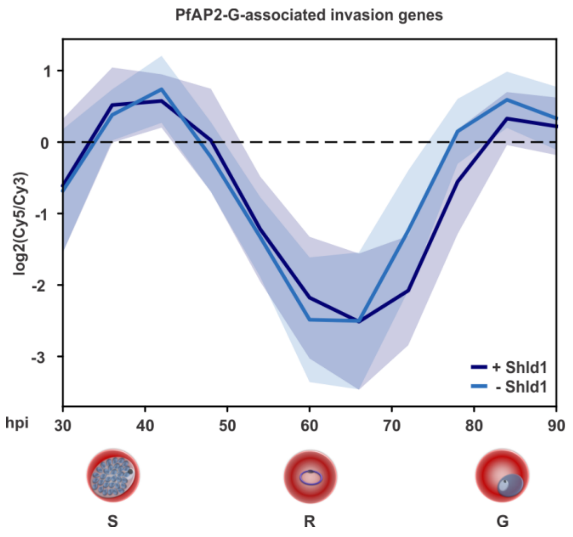


**Supplementary Figure 6:** Plot showing mean temporal expression of the 36 invasion genes bound by PfAP2-G in AP2-G-DD +Shld1 (dark blue) and AP2-G-DD -Shld1 (light blue). The shaded regions indicate the standard deviation.

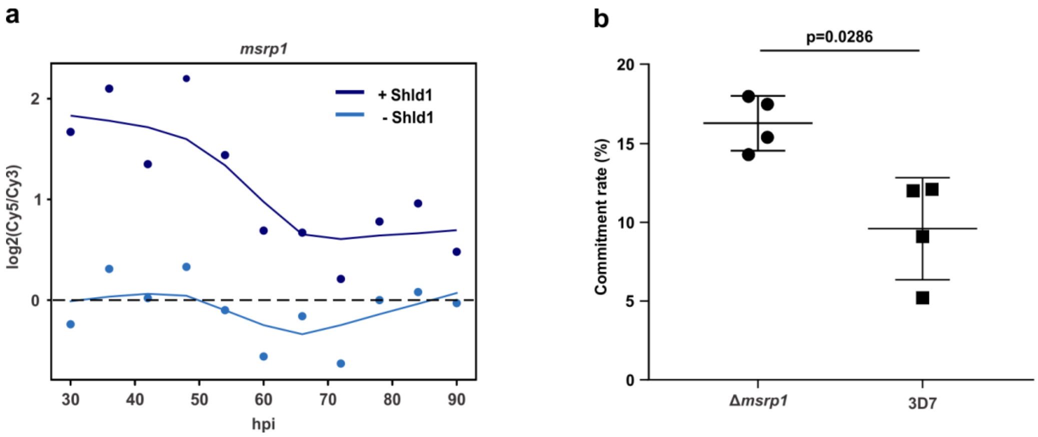


**Supplementary Figure 7: a**, Expression of *msrp1* in AP2-G-DD +Shld1 (dark blue) and AP2-G-DD -Shld1 (light blue). **b**, The *msrp1* knockout line[2] has a higher commitment rate than its parent. The horizontal bars indicate the mean and standard deviation. n = 4. The p-value was calculated using the two-tailed Mann-Whitney U test.

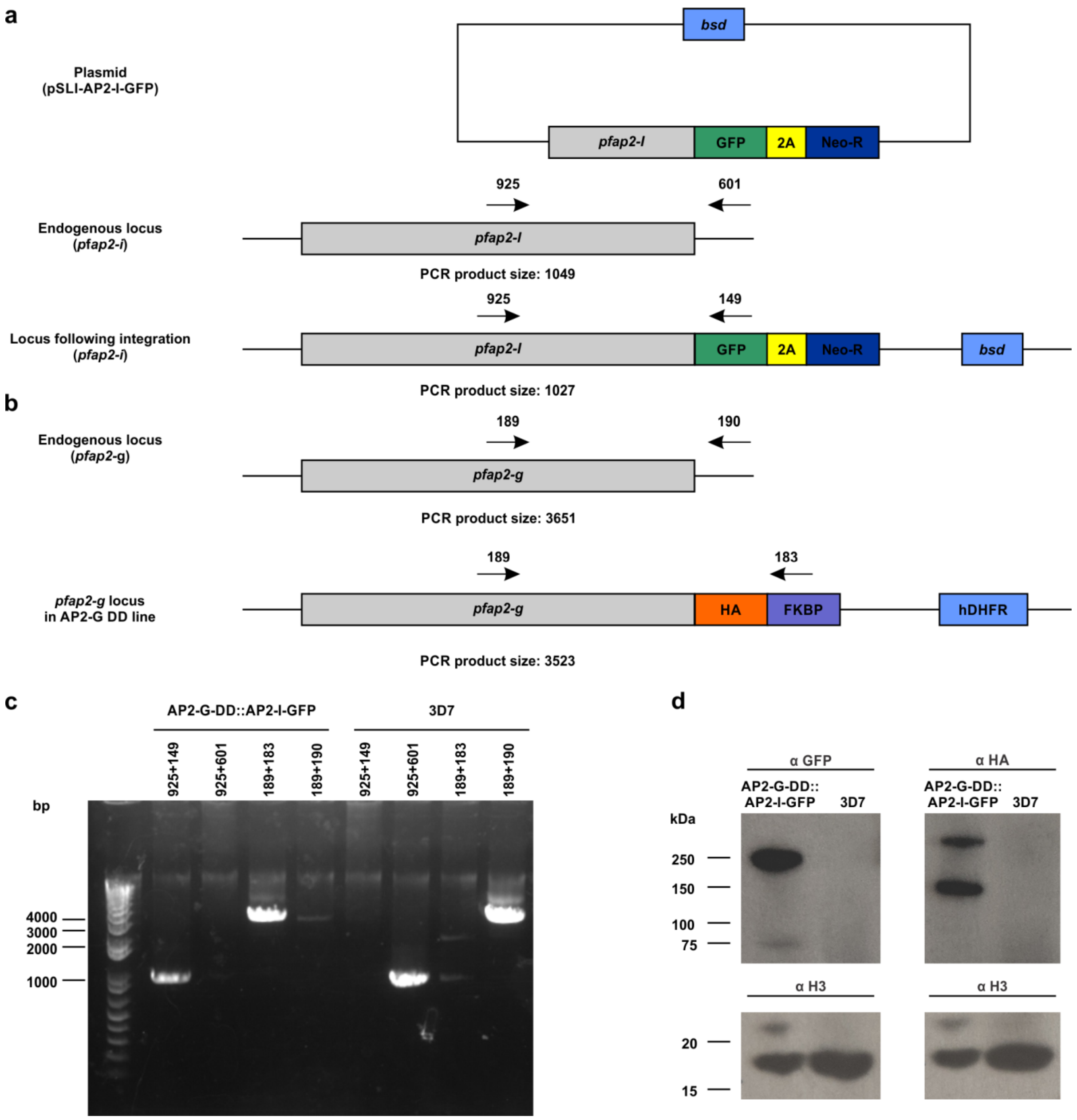


**Supplementary Figure 8: a**, The strategy used to tag PfAP2-I with GFP using selection-linked integration. Primers used to determine if integration had occurred are shown above the DNA sequences, and expected sizes of resulting PCR products are shown below. **b**, The *ap2-g* locus in wildtype and AP2-G-DD parasites. Primers used to confirm tagging of PfAP2-G are shown above the DNA sequences, and expected sizes of resulting PCR products are shown below. **c**, Genotyping PCR using the primers shown in a and b was performed on gDNA extracted from clonal AP2-G-DD::AP2-I-GFP parasites and 3D7. This confirmed that both genes had the intended integration. **d**, Western blot showing that the AP2-G-DD::AP2-I-GFP line expresses both AP2-I-GFP (top left panel) and AP2-G-DD (top right panel), but 3D7 does not. H3 (bottom panels) was used as a loading control.

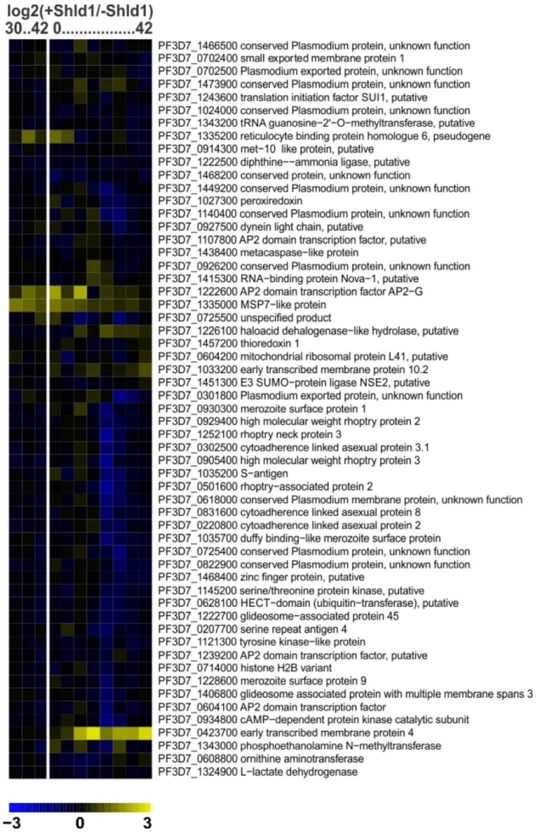


**Supplementary Figure 9:** Heatmap showing expression of genes associated with both PfAP2-G and PfAP2-I binding in AP2-G-DD +Shld1 compared to AP2-G-DD -Shld1

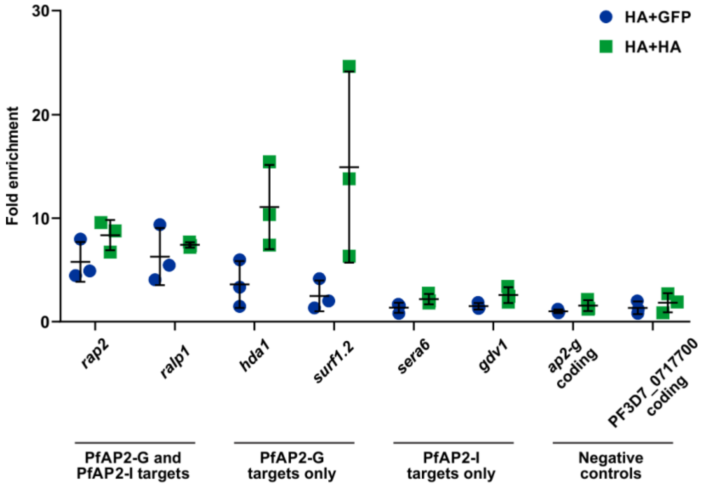


**Supplementary Figure 10:** ChIP-reChIP shows PfAP2-G and PfAP2-I bind invasion gene promoters together. Following the first round ChIP with α HA on the AP2-G-DD::AP2-I-GFP line, the immunoprecipitated material was subjected to a second of ChIP with either α GFP or α HA. Data are represented as fold enrichment relative to a negative control ChIP-reChIP with non-immune IgG. qPCR was performed to determine whether specific loci are bound simultaneously by PfAP2-G and PfAP2-I. The horizontal bars indicate the mean and standard deviation. n = 3.
